# Rapid mechanical stimulation of inner-ear hair cells by photonic pressure

**DOI:** 10.1101/2021.01.10.426123

**Authors:** Sanjeewa Abeytunge, Francesco Gianoli, A.J. Hudspeth, Andrei S. Kozlov

## Abstract

Hair cells, the receptors of the inner ear, detect sounds by transducing mechanical vibrations into electrical signals. From the top surface of each hair cell protrudes a mechanical antenna, the hair bundle, which the cell uses to detect and amplify auditory stimuli, thus sharpening frequency selectivity and providing a broad dynamic range. Current methods for mechanically stimulating hair bundles are too slow to encompass the frequency range of mammalian hearing and are plagued by inconsistencies. To overcome these challenges, we have developed a method to move individual hair bundles with photonic force. This technique uses an optical fiber whose tip is tapered to a diameter of a few micrometers and endowed with a ball lens to minimize divergence of the light beam. Here we describe the fabrication, characterization, and application of this optical system and demonstrate the rapid application of photonic force to vestibular and cochlear hair cells.

## Introduction

Hair cells in the auditory and vestibular systems of vertebrates convert mechanical stimuli into electrical signals through the process of mechanoelectrical transduction (***Hudspeth, 1989***). The mechanical receptor for such stimuli is the hair bundle, a cluster of stereocilia, or stiff enlarged microvilli, atop each hair cell. An extracellular molecular filament, the tip link, extends from the tip of each stereocilium to the side of its tallest neighbor in the plane parallel to the bundle’s axis of symmetry. Mechanically gated ion channels are located at the lower end of each tip link. When a hair bundle pivots at its base toward its tall edge in response to stimulation, the increased tension in the tip links opens the ion channels and the ensuing ionic current depolarizes the cell (Fig. 1A).

**Figure 1.**
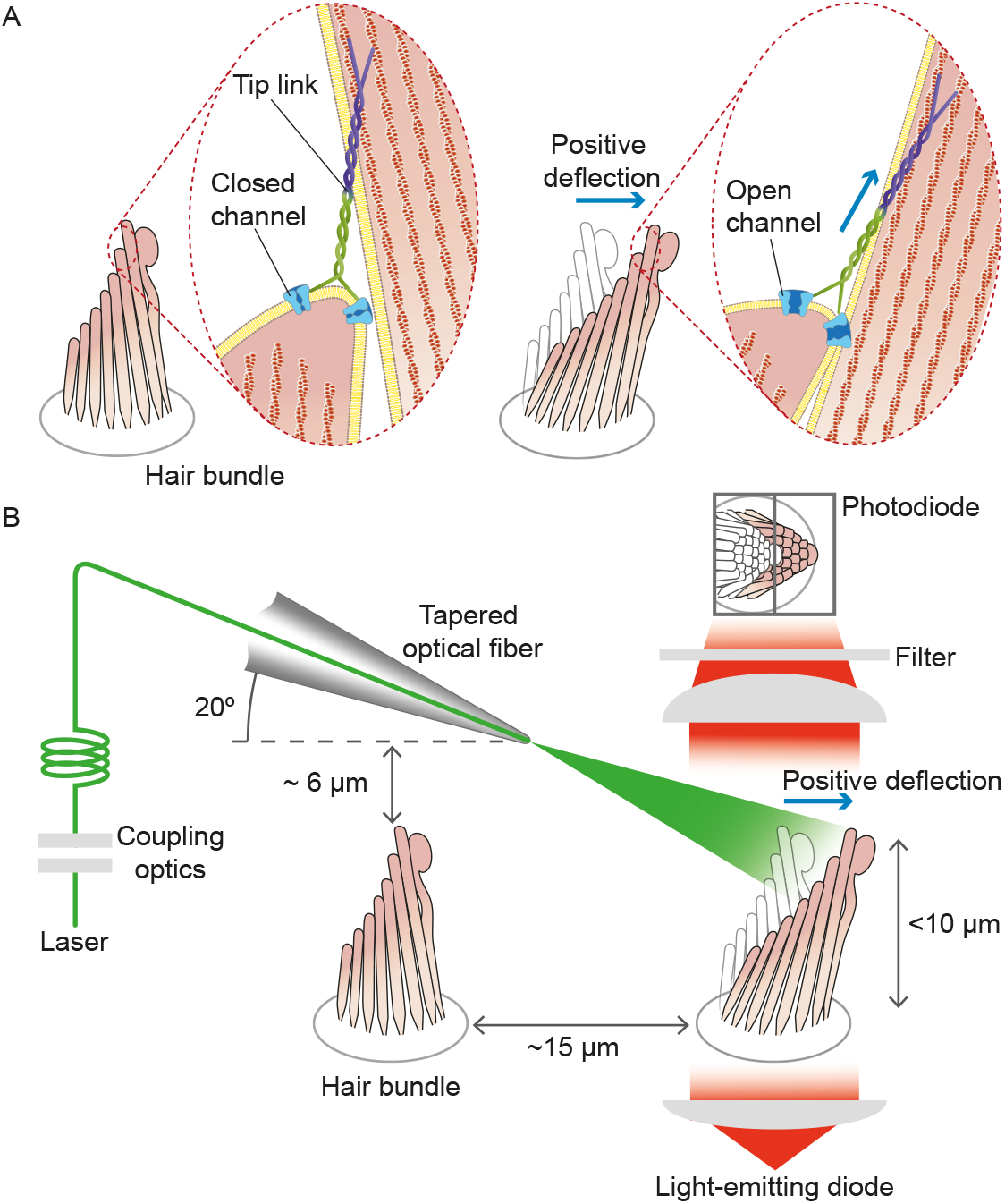
Structure of the hair bundle and configuration of the experiments. (A) A schematic illustration portrays a hair bundle, in this instance that from the bullfrog’s sacculus, at rest (left) and when deflected towards its tall edge (right). The bundle is formed by rows of stereocilia that increase in height along the axis of sensitivity and are interlinked by molecular filaments, the tip links, that stretch as the bundle moves forward. The tip links project the stimulus force onto mechanosensitive ion channels. (B) A tapered optical fiber with a spherical lens at its tip is brought within a few tens of micrometers of a hair bundle. The fiber’s angle of approximately 20° from the horizontal allows it to pass beneath the microscope’s objective lens without impinging upon other nearby hair bundles. An image of the hair bundle is projected through the microscope and onto a dual photodiode, which permits measurement of bundle motion with a precision in the nanometer range. Note that the extent of hair-bundle movement in this and the subsequent figures is greatly exaggerated for didactic purposes: the largest displacements move the bundle’s top by less than the diameter of a single stereocilium.

Although our understanding of the transduction process has improved significantly through the development of methods to mechanically stimulate a hair bundle, the techniques available nowadays pose serious limitations. Two methods are commonly used to apply force to a hair bundle. The first is to deflect the bundle with a compliant glass fiber about 100μm in length and 1 nm in diameter (***Crawford and Fettiplace, 1985**; **Howard and Ashmore, 1986**; **Howard and Hudspeth, 1988***). The fiber’s tip is attached to the top of the hair bundle and its base is driven by a piezoelectric actuator. Because the preparation is immersed in an aqueous solution, however, the fiber is subjected to hydrodynamic drag that roughly doubles that on the bundle. For a typical fiber of stiffness 500μN · m^−1^ and drag coefficient 150 nN · s · m^−1^, the time constant of responsiveness is about 300μs, which corresponds to a low-pass filter (***Crawford and Fettiplace, 1985**; **Howard and Hudspeth, 1987***) with a cutoff frequency near 500 Hz. Another problem is especially acute for the stimulation of mammalian hair bundles whose stereocilia are less cohesive than those of amphibians: when a fiber is attached at a single site in the hair bundle, the displacement of other stereocilia depends in a complex manner on elastic and hydrodynamic coupling across the bundle. This arrangement results in an uneven application of force to different stereocilia and can produce artifacts (***Indzhykulian et al., 2013**; **Nam et al., 2015***).

The second common method of stimulation uses a fluid jet that displaces a hair bundle through the action of a piezoelectric diaphragm (***Géléoc et al., 1997**; **Corns et al., 2014***). Although the resonant frequency of fluid injection can reach 5 kHz, practical use of the method is limited to less than 1 kHz owing to uncertainties in force calibration (***Dinklo et al., 2007***). Moreover, fluid leakage from the system might introduce a displacement bias.

In summary, the inability of current methods to reach higher frequencies by direct stimulation limits our quantitative understanding of hair-cell mechanics over more than 95% of the range of mammalian hearing, which extends to 20kHz in humans and at least 150kHz in some species of bats and whales. What is more, the susceptibility of current approaches to artifacts has long impeded our understanding of hearing, in particular in the case of the mammalian ear (***Nam et al., 2015***).

To address these problems, we used laser irradiation to stimulate hair bundles mechanically (Fig. 1 B). Because photonic force arises when photons are absorbed, reflected, or refracted upon interaction with an object, intense illumination should apply substantial force to a bundle. Our experiments confirmed the validity of the approach and demonstrated that the requisite irradiation does not jeopardize a bundle’s operation. This method allows us to probe hair-bundle physiology at previously inaccessible timescales, for the delivery time of the stimulus can accommodate the full frequency range of mammalian hearing. At the same time, this approach avoids the artifacts that bedevil current methods.

## Results

### Application of photonic force to a hair bundle

The conservation of momentum entails that reflected, absorbed, and refracted photons exert force on a target. All these phenomena are likely to take place when light strikes an array of stereocilia in a hair bundle. Although an analysis based on reflection alone would indicate that a hair bundle is relatively insensitive to radiation pressure, geometric considerations favor multiple modes of light propagation, each capable of transferring momentum and therefore of mechanically stimulating the bundle (see Materials and methods). Because the diameter of each stereocilium compares to the wavelength of light, the regular spacing of stereocilia within the hair bundle might additionally give rise to complex interference patterns.

#### Structure and orientation of an optical fiber

Geometrical factors are important in the stimulation of a hair bundle through an optical fiber. With its external plastic jacket, an intact fiber can be several millimeters in diameter. Even after the jacket has been removed, the core of the fiber—which is only 5 μm in diameter—lies within a cylinder of glass cladding about 125μm across. In order to bring the fiber’s core near a hair bundle without impingement of the fiber’s outer layers on the experimental preparation, it was necessary to strip the jacket and taper the cladding. By melting the tip of the fiber’s core, we created a hemispherical lens with a divergence angle in water of approximately 11° (see Materials and methods).

It was next desirable for the light beam to stimulate only a single hair bundle without affecting others nearby. This objective could be achieved readily for a flat sensory epithelium such as that of the bullfrog’s sacculus, in which the bundles are about 8 μm tall and are separated by approximately 15μm (see Fig. 1). In the rat’s cochlea, however, the distance between the row of inner hair cells and the first row of outer hair cells is only 10μm, and successive rows of outer hair cells are still more closely apposed. Moreover, this preparation is complicated by the complex curvature of its apical surface, the reticular lamina, which allows greater clearance for an optical fiber in some orientations than in others. After securing the end of a tapered optical fiber in a stable holder, we found that introducing it beneath the objective lens at an angle of 20° from the horizontal allowed the tip to approach a target hair bundle closely enough to ensure efficient stimulation, and at the same time positioned the tip far enough above other bundles to avoid damaging them (see Materials and methods).

#### Deflection of glass rods by photonic force

Before engaging in experiments with hair bundles, we conducted control experiments to confirm that photonic force from a tapered optical fiber could move an object of stiffness comparable to that of a bundle. We thinned two glass rods with a pipette puller and measured the stiffness of each by analyzing the spectrum of its Brownian motion and applying the equipartition theorem. After positioning each rod such that its shadow projected onto the photodiode, we delivered light pulses through a tapered optical fiber positioned approximately 10 μm from the rod’s tip. In both cases, we found that irradiation elicited a prompt movement in the expected direction (see ***Appendix 1*** Fig. 2). Having ascertained that our setup could deliver forces of an appropriate order of magnitude, we commenced experiments on living hair bundles.

**Figure 2.**
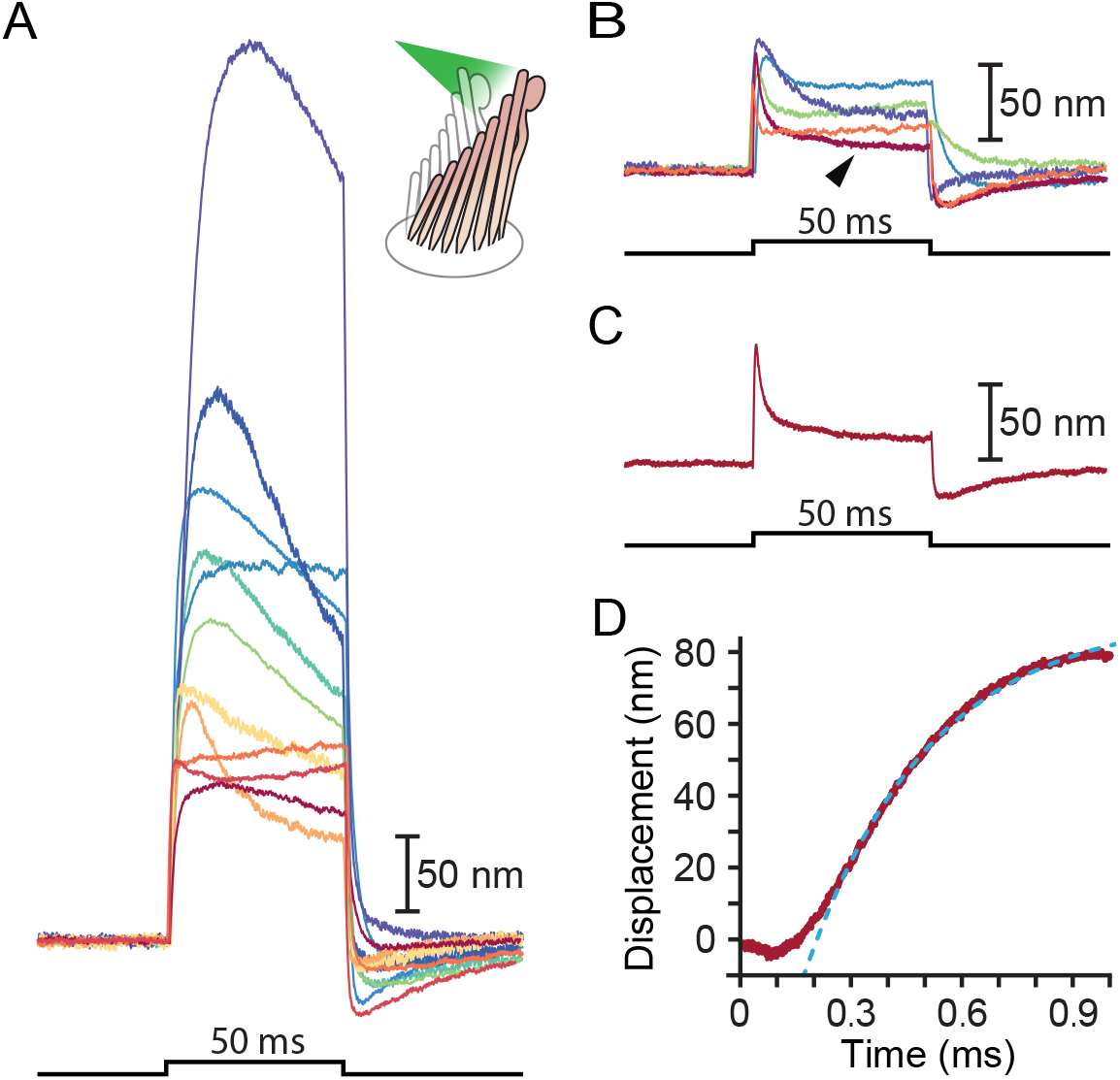
Responses of hair bundles from the bullfrog’s sacculus. (A) Although all hair bundles displayed rapid movements at the onset and conclusion of photonic stimulation, some exhibited relatively slow approaches to their peaks and slow relaxations. Eleven hair bundles were stimulated in the positive direction with 561 nm light with 30mW at the fiber’s entrance; each trace is the average of 25 responses. The schematic diagram here and in the subsequent figures shows the experimental configuration. (B) Five of the other hair bundles displayed moved rapidly at the onset of irradiation, then relaxed to plateau displacements. (C) A representative trace, marked by an arrowhead in panel B, portrays the decay of a response to a plateau level and the undershoot after stimulation characteristic of slow adaptation. (D) The rising phase of the same response is fitted with R^2^ = 0.98 to an exponential with time constant 335 μs (dashed blue line). The data at times below 250μs were not included in the fit.

### Stimulation of frog hair bundles

We stimulated 40 hair bundles of the bullfrog’s sacculus so that radiation pressure would push them toward their tall edges—the positive direction—and reliably elicited the expected movements (Fig. 2). The bundles followed similar trajectories at the onset of irradiation: the movement was approximately exponential with a time constant of (0.64 ± 0.06)ms (mean ± SEM, N = 16). The responses, which reached displacements as great as 500 nm, encompassed the range of complex trajectories reported in the literature. The relatively compliant hair bundles—those displaying initial deflections exceeding about 150nm—displayed relatively slow movements in the direction of the photonic force, a signature of the timescale of the adaptation process that allows hair cells to reset their operating points and thus detect successive stimuli (Fig. 2A) (***Ricci et al., 2000***). The “twitch,” a faster rebound of the hair bundle in a direction opposite to that of the stimulus, is another manifestation of the adaptation process that occurs instead in response to smaller movements of the hair bundle and whose magnitude decreases for larger deflections (***Ricci et al., 2000**; **Benser et al., 1996**; **Cheung and Corey, 2006***). The twitch was indeed observed in stiffer bundles with deflections of about 50 nm (Fig. 2B-D). These results indicate that photonic force is an effective means of stimulating hair bundles.

#### Polarization dependence of hair-bundle responses

Because stereocilia are densely filled with parallel actin filaments that exhibit pronounced birefringence (***Katoh et al., 1999***), we inquired how this property affected the movement of the bullfrog’s hair bundles upon photonic-force stimulation. After rupturing the tip links, we imaged a hair bundle on the dual photodiode and aligned the plane of polarization with the long axis of the stereocilia. We then rotated a half-wave plate through 90° in 10° increments. The light-induced deflection declined monotonically to an angle of 40°-50°, but remained roughly constant thereafter (see ***Appendix 1*** Fig. 3). That the response did not decline as the cosine of the angle likely reflected the fact that stereocilia are not parallel, evenly spaced cylinders but rather a more complex array with varying tilts and separations. This result nonetheless emphasized the importance of attending to the beam’s polarization, which was held parallel with the hair bundles’ long axes in subsequent experiments.

**Figure 3.**
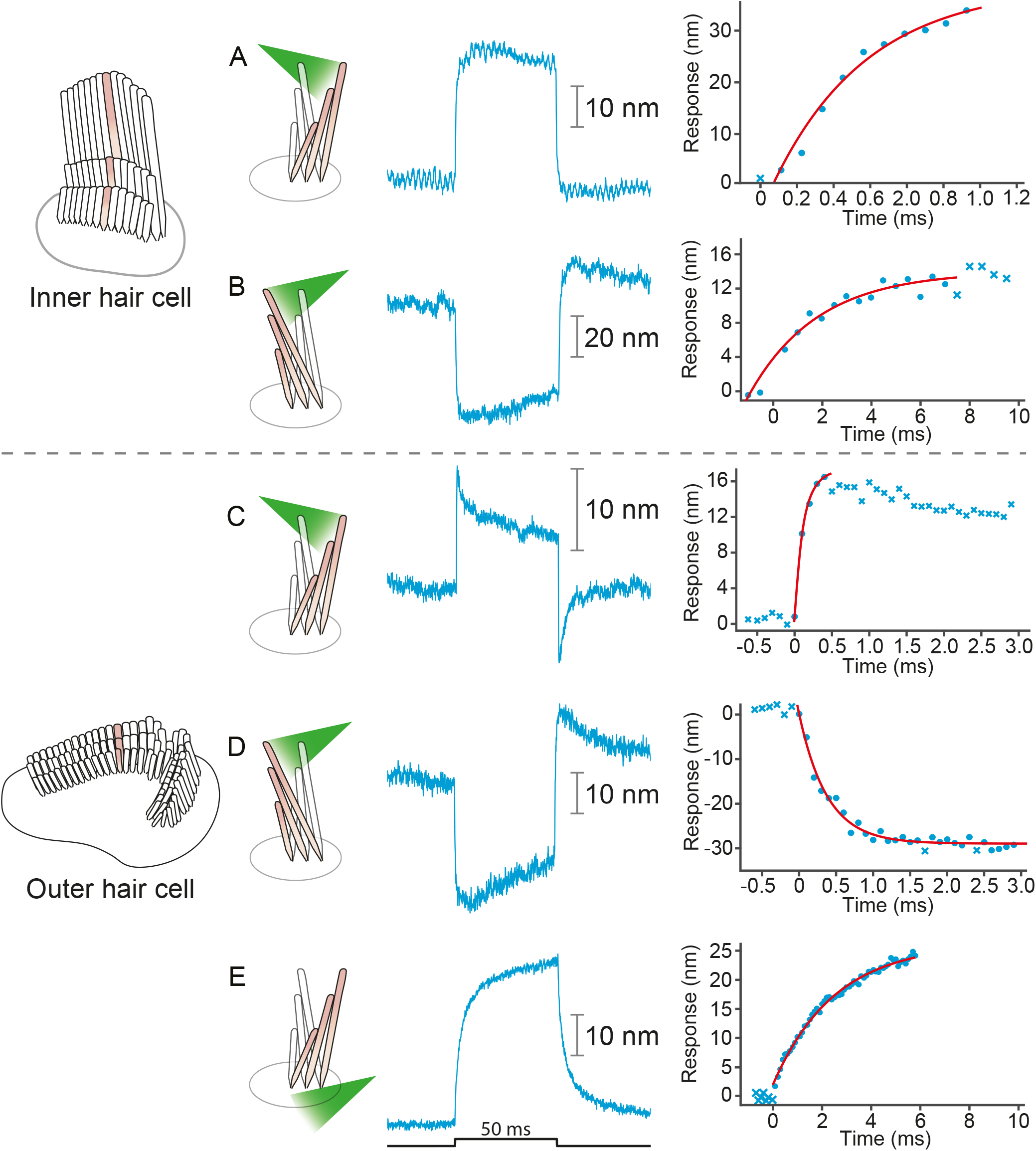
Responses of hair bundles from the rat’s cochlea. (A) Irradiation of the hair bundle from an inner hair cell evoked motion in the direction of light propagation, here the positive direction, with a time constant of 459 μs. In this and the other panels, the bundles were stimulated at 660nm with 18 mW of input power and the records represent the average of 25 repetitions. This number of repetitions was sufficient to filter the noise and isolate the characteristic shape of the hair-bundle response. (B) A similar experiment with negatively directed irradiation moved the hair bundle in the opposite direction. The time constant is 258 μs. (C) Stimulation of an outer hair cell’s bundle in the positive direction evoked a response with sharp transients at both the onset and the offset of irradiation. As shown in the associated plot, the response rose with a time constant of 123 μs and peaked in less than 1 ms. (D) Negative stimulation of an outer hair cell’s bundle evoked movement in the negative direction with an onset time constant of 377 μs. (E) When a negatively directed light beam was aimed at the soma of an outer hair cell, the bundle moved with a slow time constant of 2.1 ms in the positive direction—opposite the direction of light propagation—owing to the photothermal effect.

### Stimulation of rat hair bundles

We applied photonic stimuli to the hair bundles of both inner and outer hair cells from the cochleas of young rats. Consistent with previous evidence that mammalian hair bundles are stiffer than their amphibian counterparts (***Tobin et al., 2019***), the recorded amplitudes of deflection were typically smaller (Fig. 3A-D). The time constants for the initial displacements were again a few hundred microseconds. To characterize the efficacy of photonic stimulation for rat hair bundles, we applied positive stimuli to 22 outer hair cells from three preparations. We deflected 13 hair bundles with amplitudes varying from 25 nm to 35 nm. We also deflected seven of nine bundles from inner hair cells; the response amplitudes varied from 10nm to 75 nm and the trajectories resembled those from the frog.

### Separating the photothermal movement

As a result of localized heat generation, a hair bundle from the frog can move in the positive direction in response to laser irradiation of the cellular apex from any direction (***Azimzadeh et al., 2018***). We found that this phenomenon also occurs in hair bundles of the rat (Fig. 3E). To separate this photothermal effect from that of photonic force, we took advantage of the fact that the former requires intact tip links. When we disrupted the tip links with EDTA, we observed that both positive and negative stimuli evoked movements in the direction of light propagation (see ***Appendix 1*** Fig. 4). We also stimulated hair bundles along a direction perpendicular to their axis of symmetry and again found that they moved in the direction of photon flux (see ***Appendix 1*** Fig. 4C). These results indicate that bundle motion upon photonic stimulation can occur in the absence of a photothermal effect: bundle movements stem solely from optical radiation force.

In a frog’s hair cell, the photothermal effect apparently results from light absorption by the mitochondria that accumulate around the cuticular plate at the cell’s apical surface (***Azimzadeh et al., 2018***). Because in mammalian outer hair cells mitochondria are instead concentrated at the lateral plasma membrane (***Fuchs, 2010***), it was possible to isolate the photothermal effect by directing light well below the apical cell surface. Note that the photothermal movement was relatively slow: its time constant of 2ms was about ten times that of the movements due to photonic force. Conversely, it was possible to avert the photothermal effect by irradiating a mammalian hair bundle with intact tip links while avoiding irradiation of the cell body.

### Survival of mechanotransduction after laser irradiation

The hair bundles of healthy hair cells from the bullfrog can oscillate back-and-forth even in the absence of external stimulation (***Martin et al., 2003***). These spontaneous oscillations are a manifestation of the active process that these cells employ to amplify mechanical stimuli by counteracting viscous damping. The presence of spontaneous oscillations, which require a fully functional transduction apparatus, offers a means of assessing the viability of hair cells and the preservation of mechanotransduction following exposure to laser irradiation.

We compared the spontaneous oscillations of six hair cells before and after subjecting them to 25 pulses of light. Even at laser powers sufficient to deflect hair bundles over their entire physiological range of motion, the amplitudes and frequencies of the bundles’ oscillation were both unaffected by laser irradiation (Fig. 4). This experiment demonstrates that the mechanotransduction apparatus was not damaged by our stimulation method.

**Figure 4.**
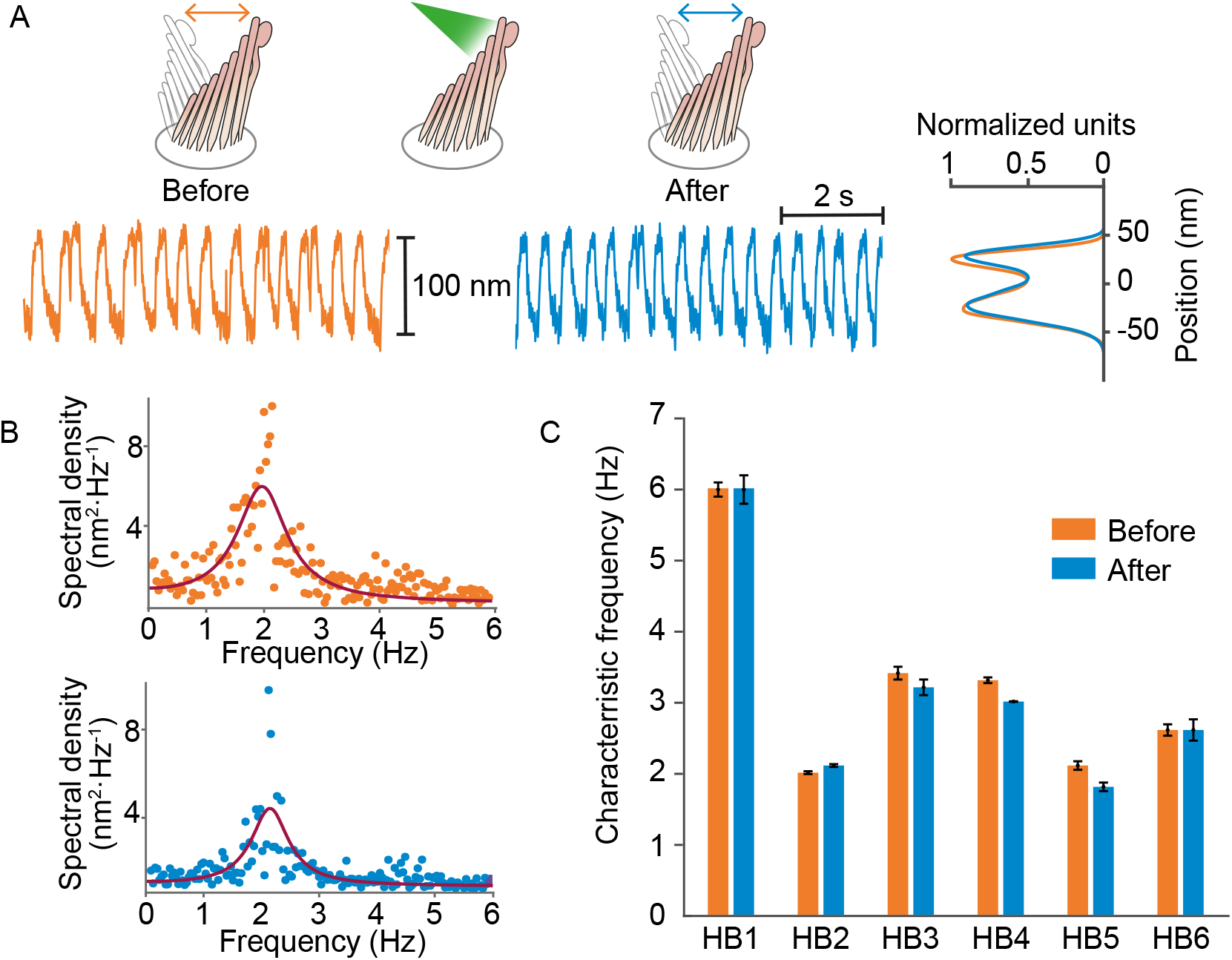
Normal operation of hair bundles after laser irradiation. (A) A hair bundle from the bullfrog displayed regular spontaneous oscillations prior to irradiation (orange). After 25 pulses of 30mW, 561 nm light from a fiber 5 mm away, the bundle continued to oscillate with a similar waveform (blue). The histograms portray the distribution of bundle positions under the two conditions and confirm that the amplitude of oscillation was similar before and after irradiation. (B) The power spectrum of the same bundle’s oscillations prior to irradiation shows a frequency peak around 2Hz, as determined by a double-Lorentzian fit. The power spectrum after irradiation has a similar peak frequency. (C) Six hair bundles (HB1 - HB6) subjected to similar treatment showed insignificant changes in their peak frequencies of oscillation after irradiation. The bundle in panels (A) and (B) is HB2.

## Discussion

We have used tapered optical fibers to apply sub-millisecond forces to individual hair bundles. The tip of each tapered fiber was small enough to be positioned near a hair bundle, and the ball lens at its tip restricted the divergence of the emitted light beam. Irradiation could be confined to a single bundle, whereupon the uniform illumination of all the stereocilia implied that each experienced a similar photonic force. It was also possible to irradiate only a portion of a bundle. Our control experiments indicated that even very extensive irradiation of the magnitude necessary to evoke large displacements did not harm the hair bundles.

The use of photonic stimulation offers at least five advantages over the currently used methods of mechanical stimulation. First, stimulation is rapid: in the present study, the rise time of mechanical responses was set by a hair bundle’s stiffness and drag coefficient, without any effect of the drag on a stimulus fiber or the inertia of a piezoelectric actuator. Second, stimulation could be made still more rapid by a process analogous to “supercharging” in a voltage-clamp system (***Armstrong and Chow, 1987***): transient irradiation with a very bright light could be used to deflect a bundle to a desired position, after which a steady force would be applied by weaker illumination during the measurement of a response. Because illumination can be switched off, a third virtue is that there is no possibility of an ill-defined steady-state offset in bundle position owing to mispositioning of a fiber or leakage from a fluid jet. The uniform illumination of the stereocilia in a bundle offers a fourth advantage, especially for mammalian hair bundles that exhibit relatively poor lateral coupling between stereocilia. And finally, photonic stimulation can be used in spaces too restricted to admit a flexible fiber or fluid jet. In particular, it should be possible to stimulate one or several hair bundles in preparations such as a hemicochlea (***He et al., 2004***) or an isolated cochlear segment (***Chan and Hudspeth, 2005a**,b*).

There are two disadvantages to photonic stimulation. Although a routine procedure after the assembly of the necessary facilities, fabrication of a tapered optical fiber requires specialized equipment and a safe environment for the use of etching solution. A second issue is calibration: unlike the force delivered by a flexible fiber, which can be calibrated through the fiber’s Brownian motion, the force exerted by photonic stimulation is not easily measured. The force can nonetheless be estimated by the use of targets whose stiffness has been independently determined, especially glass fibers such as those used in this study, or passive hair bundles including those subjected to chemical fixation.

## Materials and methods

### Estimation of photonic force

Each absorbed photon imparts all of its original momentum to the absorbing object and thereby provides an impulsive force. A reflected photon delivers twice the momentum provided by an absorbed one, whereas a refracted photon imparts momentum dependent on the angle of refraction. Because reflection sets the upper limit of the force that might be delivered to a hair bundle by a particular beam of light, we begin our analysis by treating the bundle as a perfect reflector. Averaged over one oscillation of the electromagnetic field, the radiation pressure due to illumination striking a hair bundle at an incident angle *θ* to the normal of the surface is (***Paschotta, 2010**; **Hulst, 2003***)

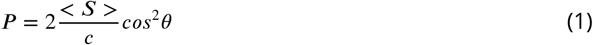

in which *P* is the radiation pressure, *S* is the average power of the electromagnetic wave, and *c* is the speed of light in vacuum. Equation 1 can alternatively be written in terms of irradiance I, or power per unit area, with units W · m^−2^, and laser power (Pwr):

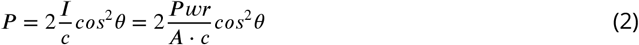

The force *F* produced at an angle *θ* is then

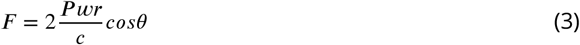

For completely absorbed photons, this relation can be modified to

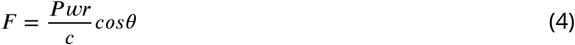

In a physiological solution, the refractive index is approximately 1.33, and therefore the speed of light is *c*/1.33. The angle of incidence in our experiments is 20°, which is set by physical clearance between the objective lens and the preparation. By Snell’s law, the angle of reflection is equal to the angle of incidence. Therefore, in the purely reflective case, using Eq. 3, we estimate that 10mW of laser power that impinges normal to the surface of the reflector generates approximately 80pN of force. However, this upper limit of the force is not achievable because stereocilia are not perfect mirrors. The actual force experienced by a hair bundle depends on the difference of the refractive indices between the solution and the stereocilia, a larger difference indicating more reflected light and larger force. As discussed below, the angle of incidence is also important.

#### Interaction of light with stereocilia: simple reflection and refraction

The interaction of light with stereocilia can be described by Fresnel equations that specify how the electric field vector’s orientation, either parallel or perpendicular to the plane of incidence, determines the amplitude of reflection and transmission (Fig. 5) (***Born et al., 1999***).

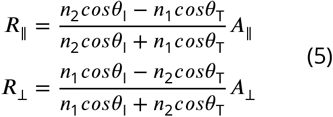

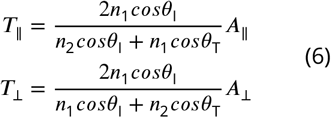

**Figure 5.**
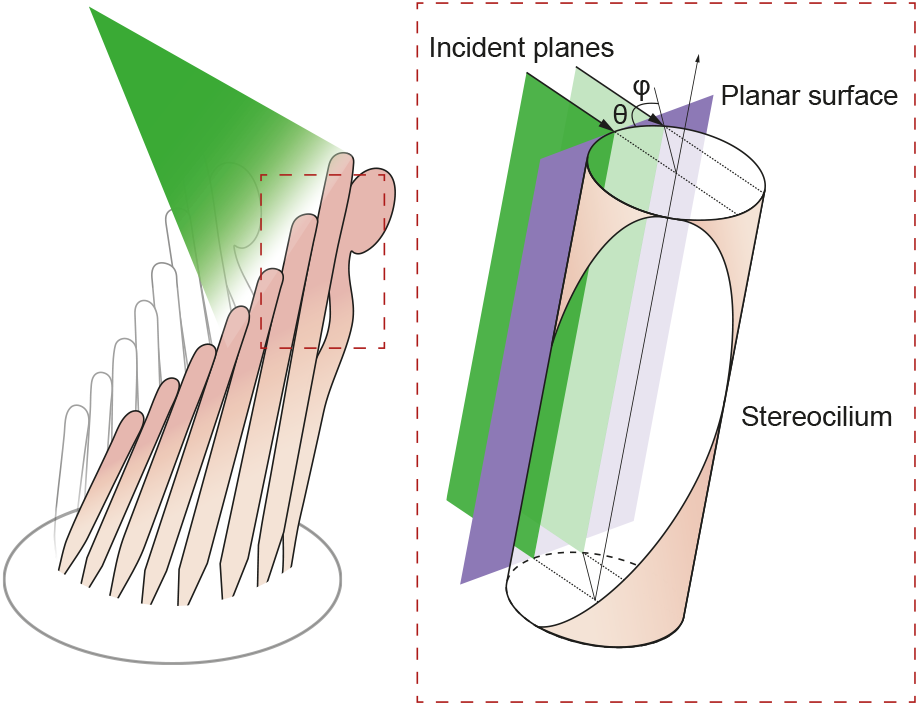
Geometry of irradiation of a cylindrical surface. As the beam from an optical fiber strikes a hair bundle, two representative sheets of the incoming light (green) are depicted along with the propagation direction of each (black arrows) and their angle of incidence *θ* with respect to the long axis of the stereocilium. Both incident planes are perpendicular to a plane tangent to the stereociliary surface (purple). Along its line of incidence onto the stereocilium, the centered sheet of light (dark green) is radially normal to the cylinder. The off-center sheet of light (pale green) strikes the stereocilium at an angle *φ* with respect to the normal.

In this set of equations, the transmission coefficient *T* or reflection coefficient *R* specify the fraction of light either reflected or transmitted at the interface of two media. The subscripts _∥_ and _⊥_ denote the orientation of the electric filed, respectively parallel or perpendicular to the plane of incidence. Light of initial amplitude *A* propagates from the medium of refractive index *η*_1_ into that of refractive index *η*_2_. The angles *θ*_1_ and *θ*_T_ are the angles of incidence and transmission (refraction), respectively. To estimate the refractive index of stereocilia we use the Gladstone-Dale relation (***Gladstone and Dale, 1863***)

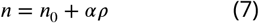

in which *η*_0_ is the refractive index of the solution, *α* is the refractive index increment for protein (***Fasman, 2020***), 200 m^3^ · kg^−1^, and *ρ* is the concentration of protein in a stereocilium, 250 kg · m^−3^ (unpublished data). We expect the refractive index of the stereocilium to be approximately 1.4. The incident angle in our apparatus is 20°, so by use of Snell’s law we find the angle of refraction for the transmitted light beam to be 19°. Applying these values to Fresnel’s equations, we calculate the following coefficients:

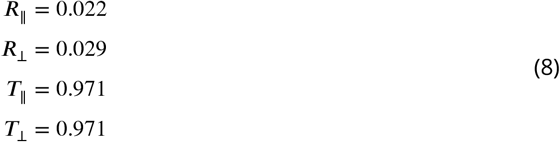

In view of the strong birefringence of stereocilia, we expect the photonic force to be greatest when the electric field is aligned parallel to a hair bundle’s vertical axis. Taking into account only the parallel components of the Fresnel equations, the coefficient of the reflected amplitude is *R*_∥_ = 0.022: approximately 0.05% of the power, or only 0.015mW of the 30mW incident on the stereocilia, should be reflected. The photonic force generated from reflection is therefore about 0.45 pN, a force unable to move a hair bundle appreciably. We must therefore reject a simple model of reflection and seek an understanding based on the reflective properties of curved surfaces.

#### Interaction of light with stereocilia: reflection from a cylindrical surface

The reflectivity, or fraction of backscattered light, is significantly higher for a curved object than for a planar one (***Ashkin, 1970***). We may analyze this effect by considering the behavior of flat sheets of light incident upon a cylinder such as a stereocilium and parallel with its long axis (Fig. 6A). Although a light sheet that strikes the stereocilium perpendicular to its surface exhibits only the effects discussed in the previous section, an off-center light sheet can produce a significantly greater force.

**Figure 6.**
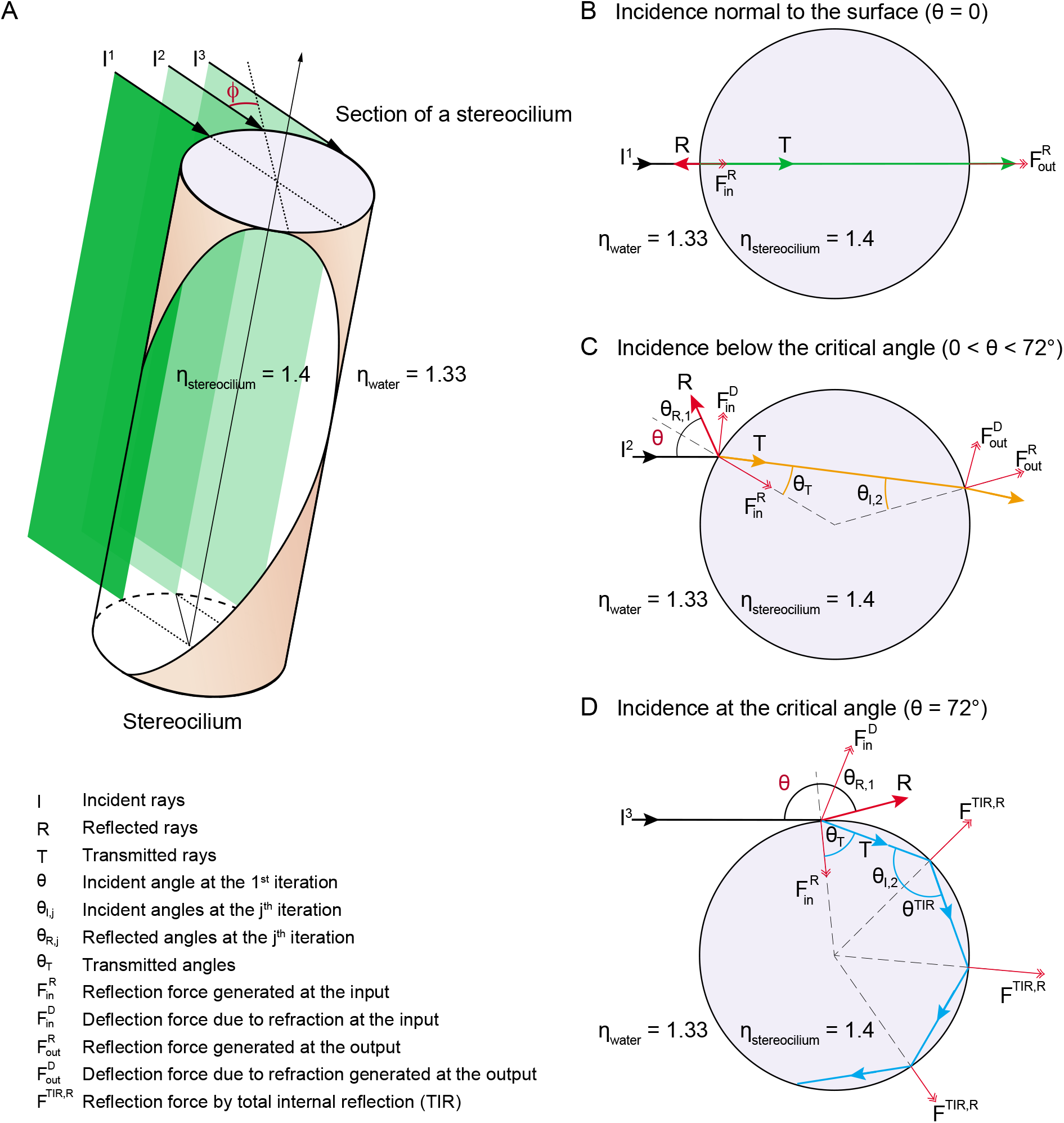
Potential fates of a plane wave incident on a stereocilium. (A) Three representative rays of a light beam interact with a stereocilium; I, R and T denote the incident, reflected, and transmitted portions of each ray. All superscripts and subscripts are defined in the figure. The rays I^1^, I^2^, I^3^ (black arrows) indicate the direction of light in water (refractive index 1.33) as it strikes a stereocilium whose refractive index is 1.4 and whose section is shown in lavender. The ray I^1^ is incident along the normal to the stereocilium, the axis of symmetry of the section. For parallel rays further from I^1^, the angle of incidence *ϕ* at which the light strikes the stereocilium’s surface increases as measured with respect to the normal. These three rays of incident light impart distinct forces on the stereocilium. (B) When a ray is reflected, it forces the stereocilium in the opposite direction and the direction of this input reflection force 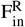 is radially aligned with the center. (C) If the ray is deflected due to refraction, a deflection force 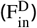 is generated on the stereocilium that is perpendicular to the direction of the ray as it propagates within the stereocilium. The incident angle is equal to the reflection angle, as is the case for the ray I^2^ as it first strikes the stereocilium (*θ* = *θ*_*R*,1_). The light that is refracted propagates along T (orange line) once inside the stereocilium until it reaches the boundary with water. At this second collision the incident angle *θ*_I,2_ is equal to the refractive angle *θ*_T_, which is too small to cause another reflection; as a result, the ray exits into water and no deflection force is generated. (D) A third kind of force arises if total internal reflection (TIR) occurs, as happens when the angle of the incident light beam is such that a ray remains trapped inside the stereocilium as it is repeatedly reflected at the boundary with water. In the case of ray I^3^, the incident angle is equal to the critical angle for total internal reflection—72° in this case—and the light remains within the stereocilium as T (blue arrow) and is reflected repeatedly each time it reaches the boundary with water. Three successive total internal reflections are shown; each generates a reflection force *F*^TIR,R^.

At any position along the stereocilium we may evaluate the behavior of representative rays of light as they impinge upon the front and back surfaces of the stereocilium. A ray exactly normal to the surface is partially reflected and partially transmitted, without refraction, through the stereocilium (Fig. 6B). This ray exerts force on the stereocilium by reflecting from its front surface, with a lesser force provided by a fraction of the transmitted light that scatters from the back surface.

A ray that strikes the stereocilium at a modest distance from its center undergoes partial reflection at the front surface, thereby producing a force in the direction of propagation and toward the stereociliary axis (Fig. 6C). Because the transmitted portion of the ray is incident upon the back surface of the stereocilium at an angle less than the critical angle for total internal reflection, it undergoes both reflection and refraction as it exits the stereocilium. That process again pushes the stereocilium in the direction of propagation as well as away from the midplane of the stereocilium.

A surprising effect ensues for a ray that impinges upon the stereocilium well away from its midplane. Such a ray undergoes partial reflection, pushing the stereocilium in the direction of propagation and toward its axis (Fig. 6D). The refracted light then strikes the back surface of the stereocilium at an angle that exceeds the critical angle for total internal reflection, which—for stereociliary cytoplasm of refractive index *η*_S_ ≈ 1.4 and water of *η*_W_ = 1.33—is approximately 72°. That ray exerts a force in the direction of light propagation and toward the stereociliary midplane. Moreover, before it eventually exits the stereocilium, the reflected ray might well undergo one or more additional total internal reflections, the first several of which exert additional force in the direction of propagation.

Because a stereocilium’s diameter is similar to the wavelength of light, its optical properties cannot be described in detail by geometric optics, but involve calculations beyond the scope of this work. However, it has been shown that for a cylinder with an aspect ratio of 15, similar to that of a stereocilium of length 8 μm and diameter 0.5 μm, the reflectivity is about 3.5 times that of a sphere of equal volume (***Gordon, 2011***). Moreover, because stereocilia are closely spaced in a regular geometric array, they likely form a grating that exhibits complex interference patterns. Nonetheless, even the qualitative description offered above emphasizes the importance of stereociliary curvature in providing exceptionally high reflection and unexpectedly great forces on stereocilia.

### Fabrication of a tapered optical fiber

In orderto produce an optical fiberwith a tip small enough to approach an individual hair bundle, it is necessary to thin the fiber’s 60 μm-thick cladding to expose the inner core of 5 μm diameter. Various methods have been employed to reduce the diameter of fibers in near-field optical microscopy and in the development of optical-fiber sensors. One common method is to use a carbon dioxide laser to machine optical fibers (***Ozcan et al., 2007***). Although this method is capable of creating symmetrical fibers, CO_2_ lasers are expensive and require complex optics. Two other methods used for removing material in optical fibers are femtosecond laser micromachining (***Wei et al., 2008a**,b; **Liao et al., 2012**; **Yuan et al., 2012***) and focused-ion-beam milling (***Kou et al., 2010**; **Yuan et al., 2011**; **André et al., 2014***). Although both methods are effective, they are time-consuming and require expensive instruments.

On the basis of previous experiments with glass fibers, we suspected that the interaction of tapered fibers with living specimens would contaminate the fibers’ tips and thus limit the use of each fiber to only a few experiments. Furthermore, the gradual degradation due to several hundred high-power optical pulses during an experiment would limit a fiber’s use to a few experiments. Both considerations required that fibers be tapered easily and cost-effectively in a typical laboratory setting. We created tapered optical fibers by Turner’s wet chemical etching with hydrofluoric acid (***André et al., 2014***). With this method, a fiber can be shaped in about 1.5 hr in any laboratory with a fume hood and few tools. In shaping each fiber, we started with a single-mode optical fiber 1 m in length and with an FC/PC connector at one end. The distal end of the fiber was prepared by stripping a 12mm length of its polymeric jacket and the polyamide coating and cleaning it with 70% ethanol. We inserted the fiber’s end into a holder that allowed it to be attached to manipulators during the fabrication process.

Etching was conducted in a fume hood (Fig. 7A). After 47.5 mL of concentrated (48%) hydrofluoric acid had been placed in a polypropylene tube (Corning Life Sciences, Tewksbury, MA, USA), 2.5 mL of red kerosene oil was added. The oil layer’s purpose was twofold. First, it provided protection to the fiber above the surface from attack by acid vapor. Second, the height of the aqueous meniscus was dependent upon the diameter of the immersed fiber, and thus declined as etching proceeded. When etching was complete, the oil layer isolated the tip from the acid.

**Figure 7.**
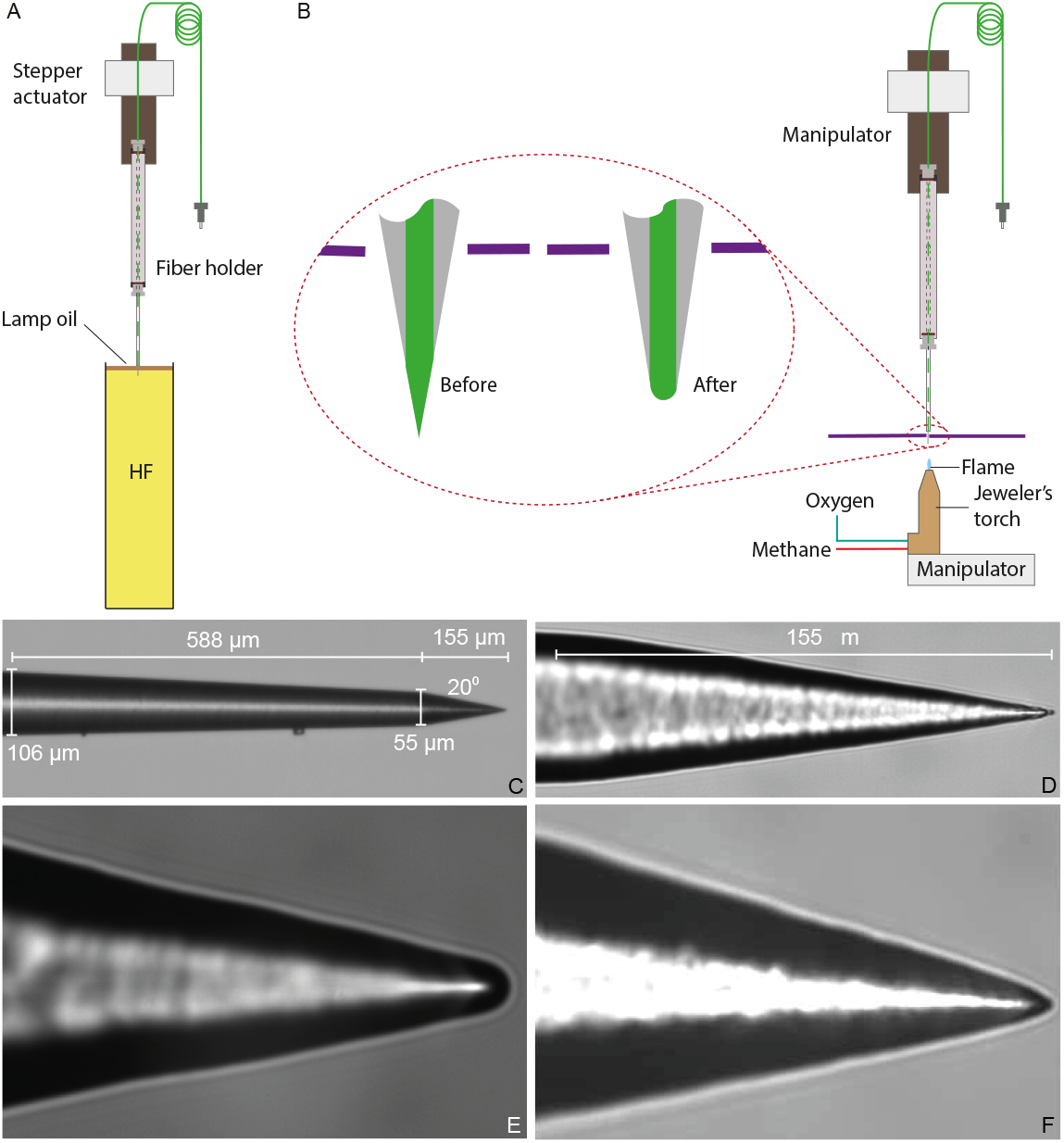
Preparation of a tapered optical fiber. (A) A schematic diagram depicts the process of reducing the optical fiber’s diameter by chemical etching. The tapered optical fiber is attached to the shaft of a linear actuator and lowered into a tube of hydrofluoric acid (HF) topped with a thin layer of lamp oil (kerosene). The convergence angle of the fiber’s tip is determined by the rate at which the tip is extracted from the acid. (B) A schematic diagram depicts the apparatus for creating a hemispherical lens at the fiber’s tip. With the aid of a three-axis micromanipulator, the fiber’s tip is inserted through a hole less than 1 mm in diameter in a horizontally mounted metal plate (purple line). The nozzle of a jeweler’s torch is aligned with the optical fiber by means of a second micromanipulator. Careful adjustment of the flow of oxygen and methane yields a flame about 0.5 mm in height. Under microscopic observation, the flame is raised until the fiber’s tip melts, whereupon the fiber is immediately retracted. (C) An image of a fiber’s tip after chemical etching with 48 % hydrofluoric acid and before polishing shows a slow taper over 588 μm followed by a steep taper over the final 155 μm. (D) At a higher magnification, the tapered but unpolished tip displays a cone angle of about 20°. (E) A high-magnification image depicts a polished tip with a relatively large hemispherical lens. (F) Another polished tip ends in a narrower lens.

The fiber’s holder was attached to a motorized linear actuator (Nanotec Electronic GmbH & Co KG, Feldkirchen, Germany) with 3 μm positioning resolution and the height of its tip was controlled through a computer interface (LabVIEW; National Instruments, Austin, TX). Because the diameter of the fiber’s tip at any point along its length depended on the duration of its immersion, it was critical to control the fiber’s extraction speed. For maximal stability during experiments, we set the length of the taper to 8 mm, the minimum required for reliable clearance of the objective lens.

After coupling a 633 nm wavelength laser to the optical fiber to render its tip visible during etching, we lowered the fiber until its tip was immediately above the interface between the oil and the acid. Under computer control, the actuator then performed a series of insertions and extractions of the fiber. The initial program inserted the fiber 10mm into the acid at 2mm · s^−1^ and extracted 8 mm at the same speed. The routine next extracted the optical fiber for 20 min at 37.5*μm* · min^−1^, reducing its diameter from 125 μm to 60 μm. The extraction then stopped and the fiber remained in the acid for 18min, during which tip was etched at a steeper angle by the gradual fall of the meniscus. The fiber was rinsed with distilled water and then with isopropyl alcohol and air-dried in the fume hood.

#### Creation of a miniature hemispherical lens

When light exits an optical fiber into a medium of lower refractive index, such as water, it diverges rapidly (***Kohls et al., 1998***). To minimize this divergence and direct the light to fall evenly upon a hair bundle, we created a focusing lens at the fiber’s tip. Although it is a common practice to attach microscopic lenses to optical fibers with flat, polished ends (***Liberale et al., 2010**; **Eversberg and Vollmann, 2015***), it was not practical do so with a taped optical fiber ending in a sharp point. We therefore created a lens by melting the fiber’s tip of silicon dioxide, which melts (***Haynes, 2011***) at 1713 °C.

We used a jeweler’s torch with a nozzle 250μm in diameter and fed with pressurized methane and oxygen. Creating a lens required clear visualization of the fiber’s sharp tip and precise manipulation of the torch (Fig. 7B). Using a pair of manual manipulators, we mounted the torch below the fiber’s tip and visualized them with a horizontal microscope. Because the hot air rising from the torch caused the thin tip of the fiber to flutter, we reduced the convection around the fiber by partially exposing the tip to the torch’s flame through an aperture 1 mm in diameter in a square metal plate 50mm on each side. After focusing the image of the fiber’s tip on an eyepiece reticle, we carefully raised the unlit torch toward the fiber and aligned the two to prevent asymmetry in the lens. The torch was then lowered, lit, and adjusted to a flame height of about 0.5 mm. As we then raised the torch, the core of the fiber began to melt and promptly assumed a hemispherical shape (Fig. 7C-F). We immediately lowered the torch and allowed the fiber to cool before removing it from the apparatus.

#### Estimation of the area of irradiation

After fabricating a tapered optical fiber of suitable shape, we characterized its pattern of illumination before using it in experiments. This process was designed to evaluate the optimal distance between the fibers’ tip and a hair bundle so that we could match the diameter of the light spot to the bundle’s width.

After coupling a laser to the tapered fiber, we approximated the fiber’s tip to the flat end of an ordered fiber bundle and monitored their separation under a microscope with a camera (Fig. 8A). Passing through a droplet of water, the light from the tapered fiber impinged on the fiber cores of the bundle and propagated to the distal end, where it was imaged through a dry objective lens (Plan 10X, numerical aperture 0.25, Olympus, Tokyo, Japan) onto second camera. The illuminated fiber cores defined the diameter of the illuminated area on the fiber bundle (Fig. 8B). By capturing images at intervals of 100μm as the tip of the tapered fiber was withdrawn from the fiber bundle, and measuring the diameter of the illuminated area at 95 % of the power spread, we estimated the divergence angle of the light cone.

**Figure 8.**
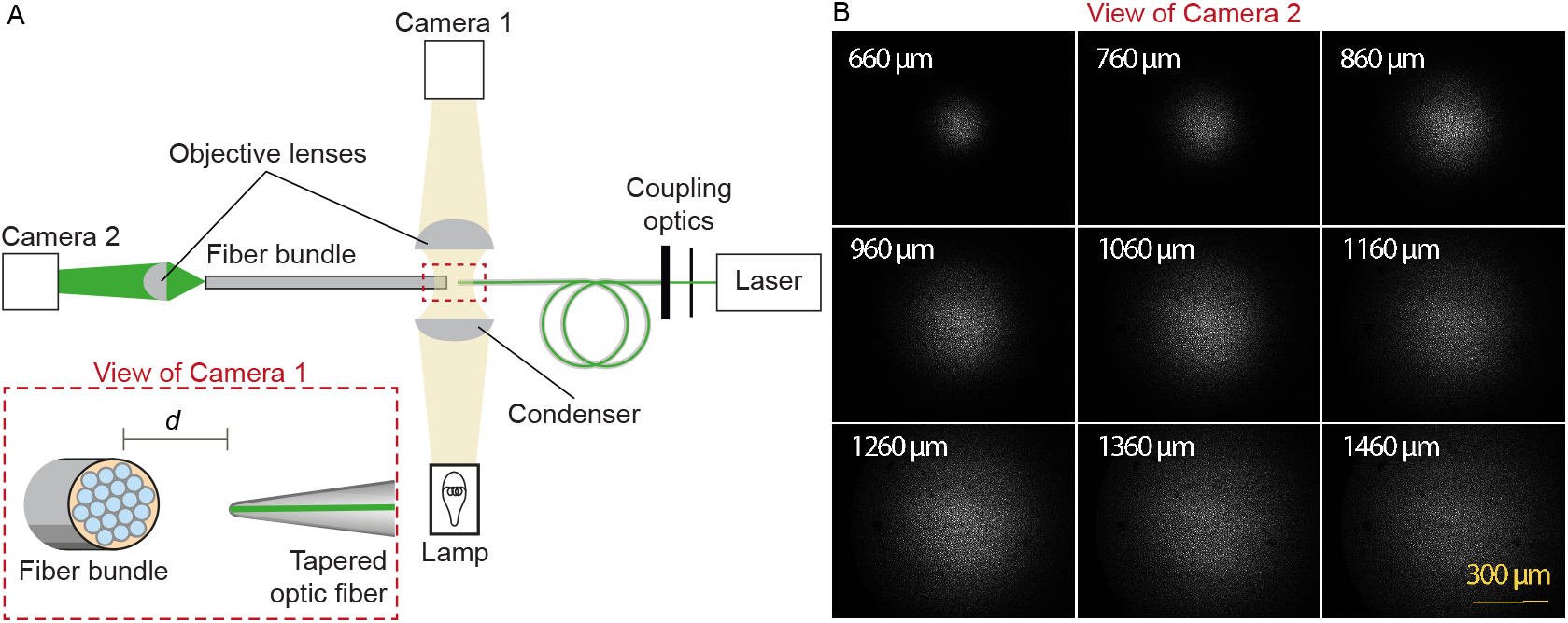
Characterization of the light spot produced by a polished fiber. (A) A schematic diagram depicts the apparatus for characterization of a fiber’s output. During observation by camera 1 attached to the microscope, the tapered optical fiber is coupled to the laser and brought near the transverse surface of a fiber bundle (IGN-8/30, Sumitomo Electric, Japan). The view through camera 1, with the fiber tip pointing at one end of the fiber bundle a distance d away, is schematized in the inset. The fiber bundle has 30,000 inner cores, each 2 μm in diameter and with a center-to-center spacing of 4 μm. The distal end of the bundle is imaged by camera 2 at a magnification of 11.6X. (B) Images of the output, as captured with camera 2, show the divergence of the light beam as the tapered optical fiber is brought approximately 660 μm from the bundle and retracted by intervals of 100μm.

### Experimental configuration

During each experiment, the tapered optical fiber was inserted through a glass capillary placed in a custom-made electrode holder that could be affixed to a micromanipulator (Fig. 9A). This holder ensured that the fiber’s tip was stable despite possible vibrations or displacements of the remainder of the fiber.

**Figure 9.**
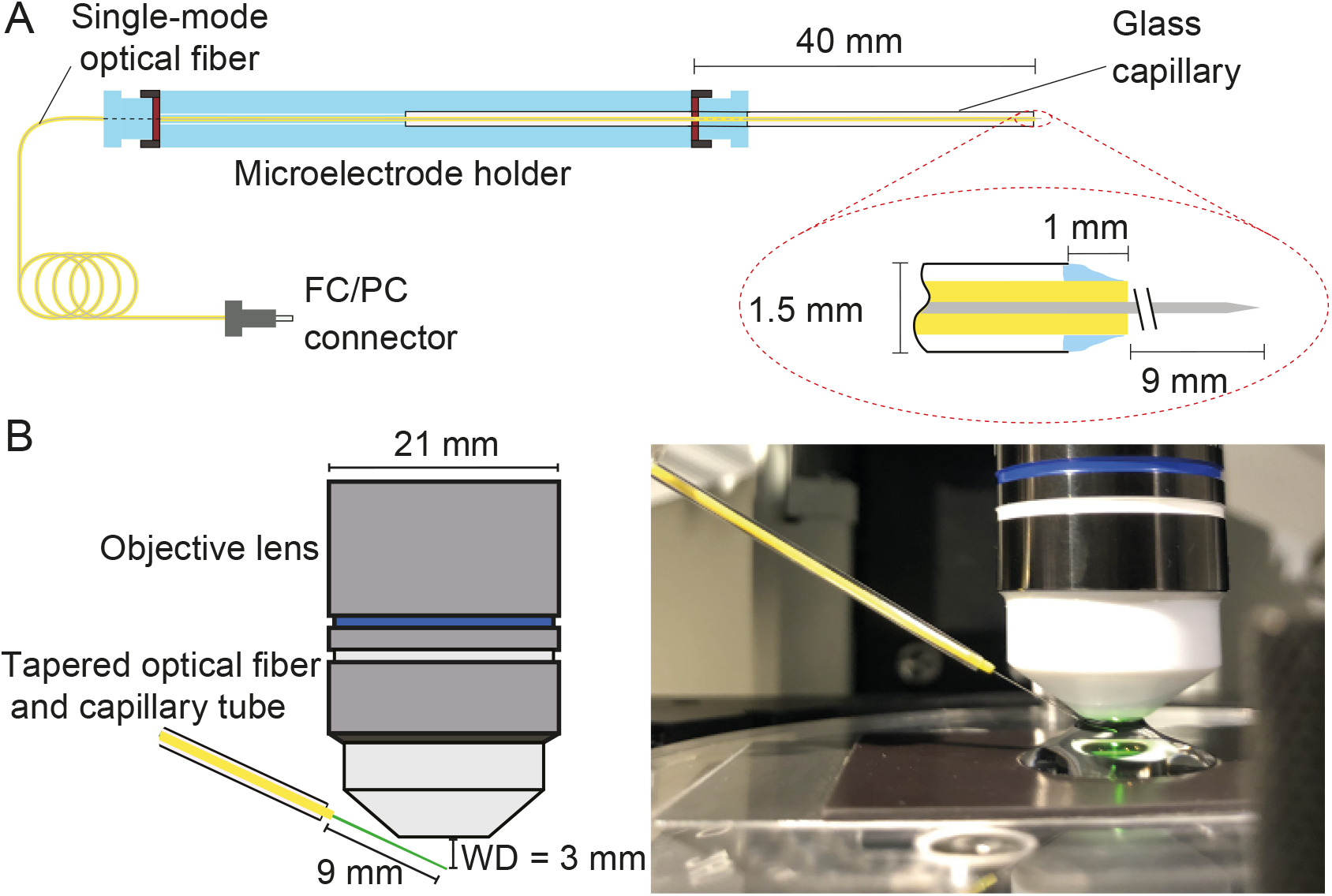
Positioning fiber under an objective lens using a custom-made holder. (A) Holder for the tapered optical fiber. The schematic drawing portrays a tapered optical fiber inserted in the mount constructed from a microelectrode holder and a glass capillary, from which the fiber’s distal tip protrudes 10mm. The coiled optical fiber’s inner core is depicted in gray inside the yellow jacket. As seen in the inset, the space between the glass capillary and the yellow jacket that protrudes 1 mm past the capillary’s tip is packed with vacuum grease (light blue). The distal end of the fiber is terminated with an FC/PC connector.Positioning of a fiber under an objective lens. (B) A schematic drawing (left) shows the length of the glass capillary that protrudes from the fiber-holder relative to the objective lens. The photograph (right) shows the tapered optical fiber, objective lens of the microscope, and preparation chamber in an experiment.

Under the control of a micromanipulator (ROE 200, Sutter Instruments, Novato, CA, USA), the fiber’s distal end was introduced into the experimental chamber beneath a 60X water-immersion objective lens (LUMPlanFL N, numerical aperture 1.0, Olympus, Tokyo,Japan). The incidence angle of about 20° with respect to the horizontal ensured that the fiber cleared both the upper edge of the experimental chamber and the lower rim of the lens (Fig. 9B).

Light from a 600 nm light-emitting diode (Prizmatix Ltd., Southfield, MI, USA) illuminated the specimen through an inverted 60X water-immersion objective lens (LUMPlan FI/IR, numerical aperture 0.9, Olympus) that served as a condenser (see ***Appendix 1*** Fig. 1). To permit differential-interference imaging, a polarizer was positioned just above the microscope’s field diaphragm and a crossed analyzer above its tube lens, and both objective lenses were equipped with Wollaston prisms. A rotating quarter-wave plate above the polarizer permitted optimization of the image, and a heat filter protected the specimen from infrared damage. Light that had traversed the specimen, the objective lens, and the tube lens was relayed by two mirrors and projected with a total magnification of 900X onto a dual photodiode, which permitted measurements of hair-bundle movement with nanometer precision. A dichroic mirror imposed before the photodiode prevented contamination of the movement signal by light from the stimulating laser. For the selection of appropriate hair bundles, the light path could be diverted to a camera that permitted observation of the specimen on a digital monitor.

## Acknowledgments

We thank Brian Fabella for assistance with the fabrication of apparatus and the programming of data-acquisition software and Rodrigo Alonso for the preparation of bullfrog sacculi. The members of our research groups graciously provided critical comments on the manuscript. A.J.H. is an Investigator of Howard Hughes Medical Institute. A.S.K. was supported by grants 108034/Z/15/Z and 214234/Z/18/Z from the Wellcome Trust and by an Imperial College Network of Excellence Award.

## Appendix 1

**Appendix 1 Figure 1.**
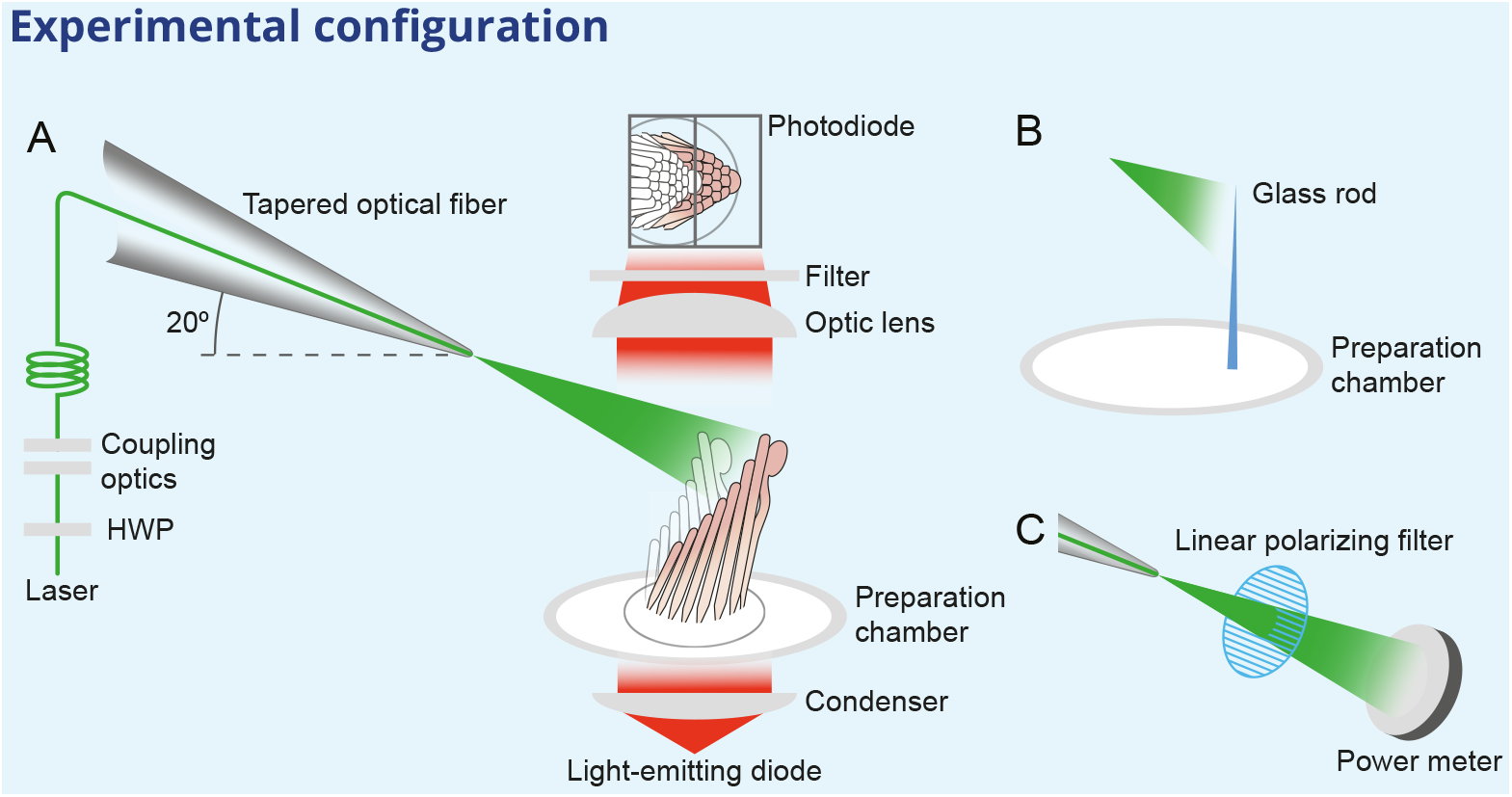
Experimental configuration. The schematic drawing shows the arrangement of the main components in the experimental apparatus. (A) The laser beam of 561 nm wavelength (green line) traverses the half-wave plate (HWP) and is coupled to the tapered optical fiber. The fiber’s distal end with the microlens is approximated to the experimental preparation under a microscope. Illumination from a light-emitting diode with a central wavelength of 660nm (red) is focused by a condenser onto a hair bundle, which is imaged onto a dual photodiode with a 60X objective lens of numerical aperture of 1.0. A dichroic mirror blocks 562nm laser light while it passes the light coming from the light-emitting diode. (B) In control experiments, a glass rod (blue triangle) of stiffness comparable to that of a bullfrog’s hair bundle is mounted vertically in the experimental chamber. The laser light (green triangle) deflects the rod, whose motion can be measured with the system in panel A. (C) To orient the plane of polarization of light with the long axis of the stereocilia, light from the tapered optical fiber is passed through a linear polarizing filter and its intensity is measured with a power meter. The maximal power is detected when the polarization plane is aligned with the transmission direction of the polarizing filter.

**Appendix 1 Figure 2.**
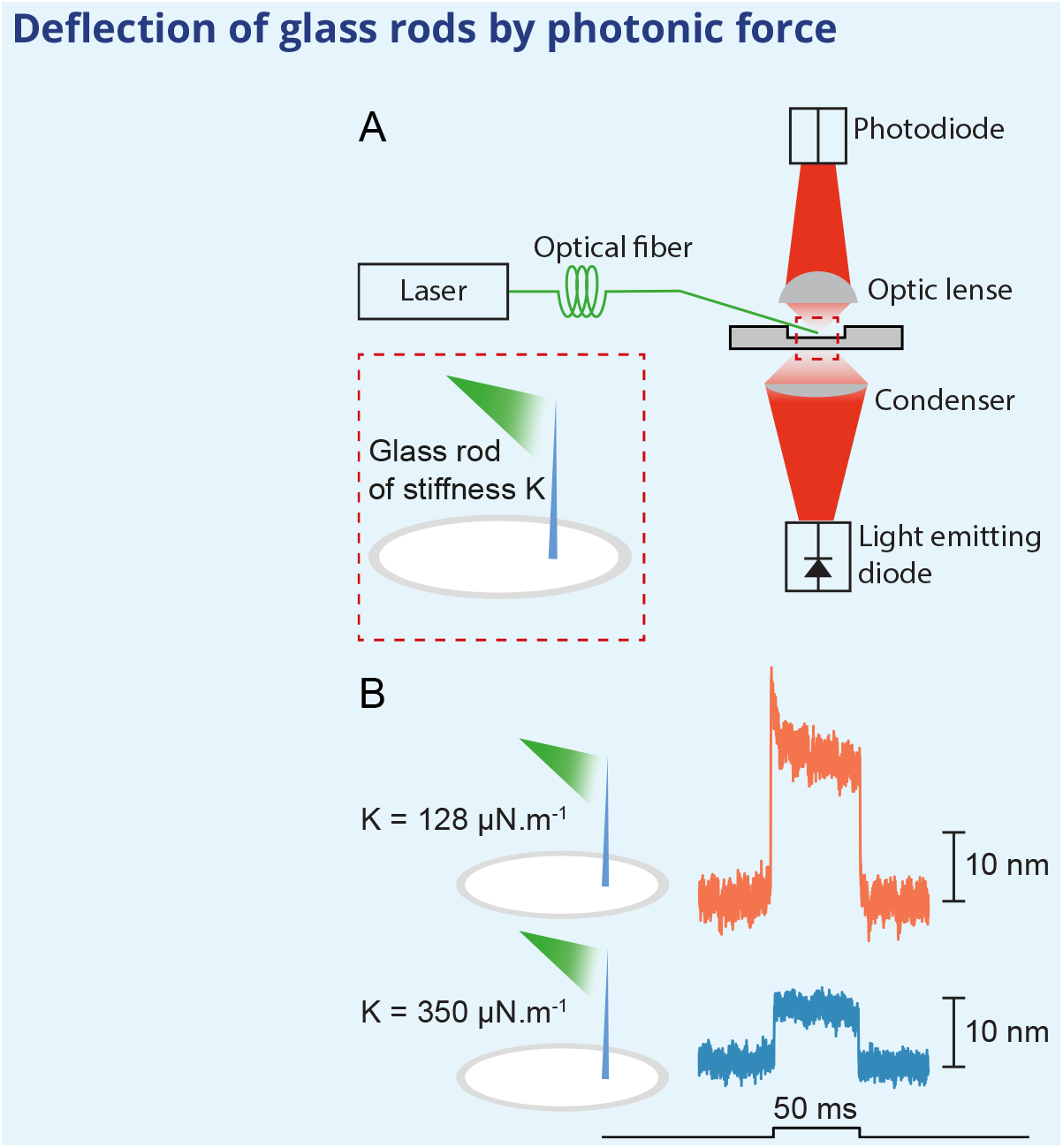
Application of photonic force applied to glass rods. (A) Two glass rods were placed in the experimental chamber and irradiated through a tapered optical fiber for 50ms. The average of 25 deflections was recorded for each rod. (B) The glass rod with lower stiffness of 128 μN · m^−1^ (orange) moved thrice as far as the fiber with a higher stiffness of 350μN · m^−1^ (blue). The estimated power of irradiation falling upon each rod was 20mW at a wavelength of 561 nm. The sudden movement at the onset of illumination for the rod of lower stiffness likely stemmed from thermoelastic effects.

**Appendix 1 Figure 3.**
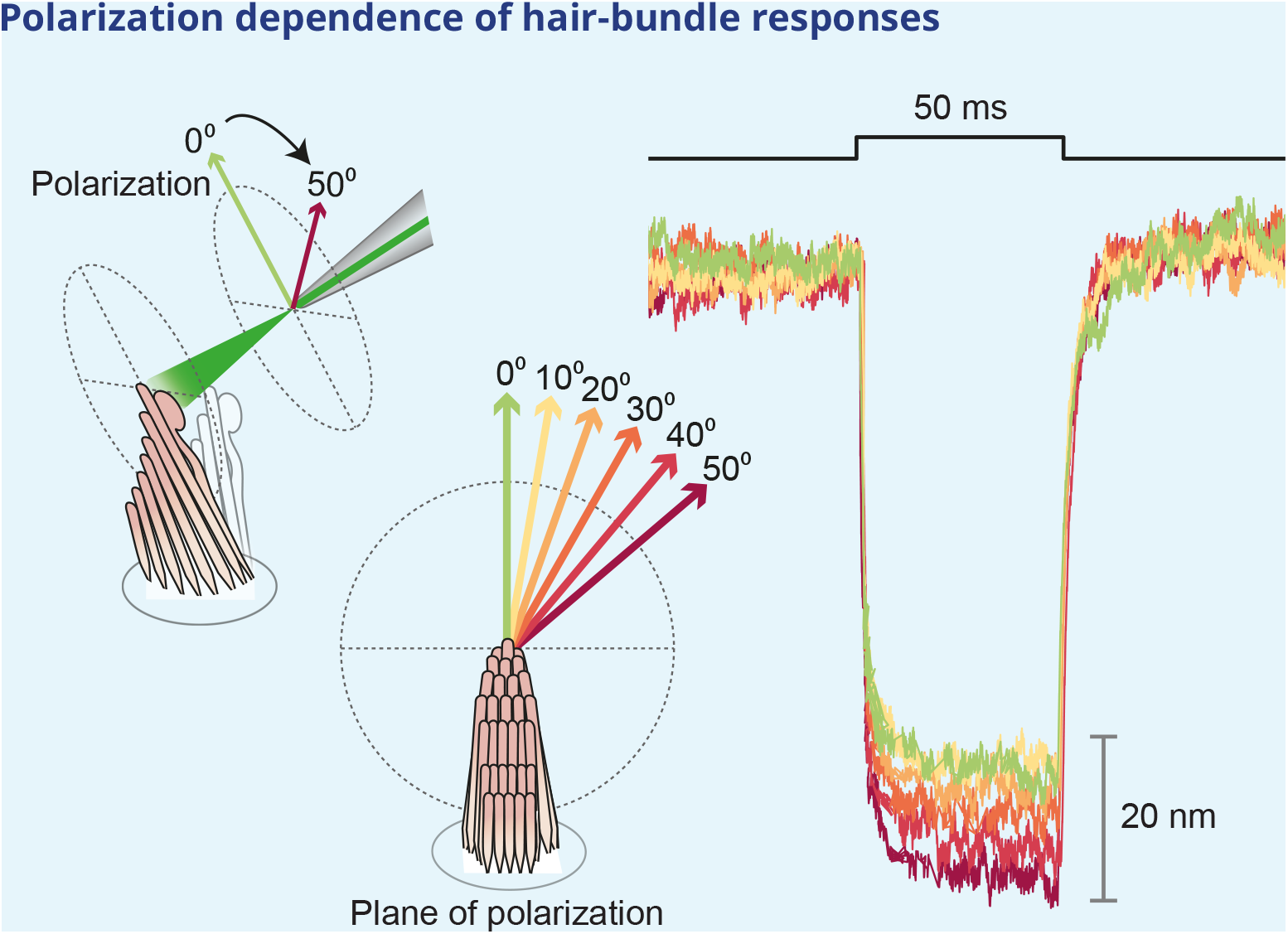
Effect of polarization on response amplitude. After its tip links had been broken by exposure to 5 mM BAPTA for 30 s, a hair bundle from the bullfrog’s sacculus was stimulated in the negative direction with 50ms, 30mW laser pulses. Using a half-wave plate between the laser and the coupling optics, we rotated the polarization plane about the axis of propagation between 0° and 50°. For a simple polarized object, the reflected power should decline by the cosine of the angle. Although the results showed a qualitative agreement with the prediction, we observed a significantly smaller reduction in amplitude consistent with the fact that stereocilia are birefringent, but exhibit significant scattering of light at all angles. Each trace represents the average of 25 recordings.

**Appendix 1 Figure 4.**
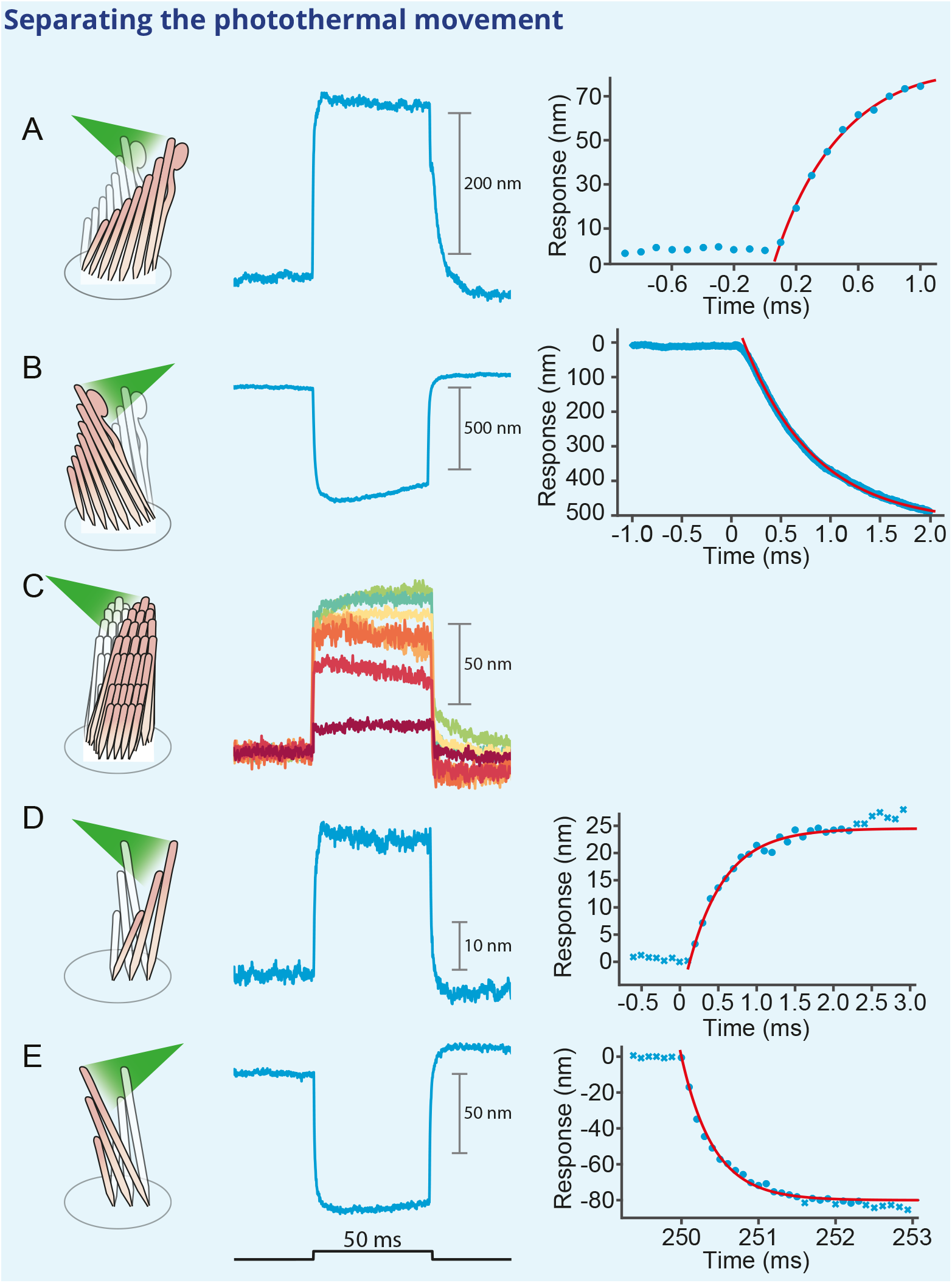
Deflection of hair bundles by optical radiation force without a photothermal effect. (A) After tip links had been ruptured by a Ca^2+^ chelator, photonic force displaced a bullfrog’s bundle in the positive direction with a time constant of 415μs. In this and the other panels, the bundles were stimulated at 561 nm with 30mW of input power and the records represent the average of 25 repetitions. (B) Stimulation in the negative direction evoked a negative movement with a time constant of 750μs. (C) Photonic force applied at 90° to the axis of sensitivity displaced a hair bundle in the direction of irradiation. (D) After the disruption of tip links, the hair bundle from a rat’s outer hair cell moved with a time constant of 467 μs in the direction of photonic stimulation. (E) Negatively directed stimulation conversely evoked motion with a time constant of 418 μs in the negative direction.

## References

André RM, Pevec S, Becker M, Dellith J, Rothhardt M, Marques MB, Donlagic D, Bartelt H, Frazão O. Focused ion beam post-processing of optical fiber Fabry-Perot cavities for sensing applications. Optics Express. 2014 Jun; 22(11):13102–13108. https://www.osapublishing.org/oe/abstract.cfm?uri=oe-22-11-13102, doi: 10.1364/OE.22.013102.

Armstrong CM, Chow RH. Supercharging: a method for improving patch-clamp performance. Biophysical Journal. 1987 Jul; 52(1):133–136.

Ashkin A. Acceleration and Trapping of Particles by Radiation Pressure. Physical Review Letters. 1970 Jan; 24(4):156–159. https://link.aps.org/doi/10.1103/PhysRevLett.24.156, doi: 10.1103/PhysRevLett.24.156.

Azimzadeh JB, Fabella BA, Kastan NR, Hudspeth AJ. Thermal Excitation of the Mechanotransduction Apparatus of Hair Cells. Neuron. 2018; 97(3):586–595.e4. doi: 10.1016/j.neuron.2018.01.013.

Benser ME, Marquis RE, Hudspeth AJ. Rapid, active hair bundle movements in hair cells from the bullfrog’s sacculus. The Journal of neuroscience: the official journal of the Society for Neuroscience. 1996 Sep; 16(18):5629–5643.

Born M, Wolf E, Bhatia AB, Clemmow PC, Gabor D, Stokes AR, Taylor AM, Wayman PA, Wilcock WL. Principles of Optics: Electromagnetic Theory of Propagation, Interference and Diffraction of Light. 7th edition ed. Cambridge; New York: Cambridge University Press; 1999.

Chan DK, Hudspeth AJ. Ca2+ current-driven nonlinear amplification by the mammalian cochlea in vitro. Nature neuroscience. 2005 Feb; 8(2):149–155. doi: 10.1038/nn1385.

Chan DK, Hudspeth AJ. Mechanical responses of the organ of corti to acoustic and electrical stimulation in vitro. Biophysical journal. 2005 Dec; 89(6):4382–4395. doi: 10.1529/biophysj.105.070474.

Cheung ELM, Corey DP. Ca2+ changes the force sensitivity of the hair-cell transduction channel. Biophysical journal. 2006 Jan; 90(1):124–139. doi: 10.1529/biophysj.105.061226.

Corns LF, Johnson SL, Kros CJ, Marcotti W. Calcium entry into stereocilia drives adaptation of the mechanoelectrical transducer current of mammalian cochlear hair cells. Proceedings of the National Academy of Sciences of the United States of America. 2014 Sep; doi: 10.1073/pnas.1409920111.

Crawford AC, Fettiplace R. The mechanical properties of ciliary bundles of turtle cochlear hair cells. The Journal of physiology. 1985 Jul; 364:359–379.

Dinklo T, Meulenberg CJW, van Netten SM. Frequency-dependent properties of a fluid jet stimulus: calibration, modeling, and application to cochlear hair cell bundles. Journal of the Association for Research in Otolaryngology: JARO. 2007 Jun; 8(2):167–182. doi: 10.1007/s10162-007-0080-0.

Eversberg T, Vollmann K. Spectroscopic Instrumentation: Fundamentals and Guidelines for Astronomers. Astronomy and Planetary Sciences, Berlin Heidelberg: Springer-Verlag; 2015. https://www.springer.com/gp/book/9783662445341, doi: 10.1007/978-3-662-44535-8.

Fasman GD. Handbook of Biochemistry: Section A Proteins, Volume II. Routledge & CRC Press; 2020. https://www.routledge.com/Handbook-of-Biochemistry-Section-A-Proteins-Volume-II/Fasman/p/book/9781315893303.

Fuchs PA, editor. Oxford Handbook of Auditory Science: The Ear. Oxford University Press; 2010. https://www.oxfordhandbooks.com/view/10.1093/oxfordhb/9780199233397.001.0001/oxfordhb-9780199233397, doi: 10.1093/oxfordhb/9780199233397.001.0001.

Gladstone JH, Dale TP. XIV. Researches on the refraction, dispersion, and sensitiveness of liquids. Philosophical Transactions of the Royal Society of London. 1863 Jan; 153:317–343. https://royalsocietypublishing.org/doi/abs/10.1098/rstl.1863.0014, doi: 10.1098/rstl.1863.0014.

Gordon HR. Light scattering and absorption by randomly-oriented cylinders: dependence on aspect ratio for refractive indices applicable for marine particles. Optics Express. 2011 Feb; 19(5):4673–4691. https://www.osapublishing.org/oe/abstract.cfm?uri=oe-19-5-4673, doi: 10.1364/OE.19.004673.

Géléoc GS, Lennan GW, Richardson GP, Kros CJ. Aquantitative comparison of mechanoelectrical transduction in vestibular and auditory hair cells of neonatal mice. Proceedings Biological Sciences. 1997 Apr; 264(1381):611–621. doi: 10.1098/rspb.1997.0087.

Haynes WM, editor. CRC Handbook of Chemistry and Physics, 92nd Edition. 92nd edition ed. Boca Raton, Fla.: CRC Press; 2011.

He DZZ, Jia S, Dallos P. Mechanoelectrical transduction of adult outer hair cells studied in a gerbil hemicochlea. Nature. 2004 Jun; 429(6993):766–770. http://www.nature.com/doifinder/10.1038/nature02591, doi: 10.1038/nature02591.

Howard J, Ashmore JF. Stiffness of sensory hair bundles in the sacculus of the frog. Hearing research. 1986; 23(1):93–104.

Howard J, Hudspeth AJ. Mechanical relaxation of the hair bundle mediates adaptation in mechanoelectrical transduction by the bullfrog’s saccular hair cell. Proceedings of the National Academy of Sciences of the United States of America. 1987 May; 84(9):3064–3068.

Howard J, Hudspeth AJ. Compliance of the hair bundle associated with gating of mechanoelectrical transduction channels in the bullfrog’s saccular hair cell. Neuron. 1988 May; 1(3):189–199.

Hudspeth AJ. How the ear’s works work. Nature. 1989 Oct; 341(6241):397–404. doi: 10.1038/341397a0.

Hulst HCvd. Light Scattering by Small Particles. Illustrated edition ed. New York: Dover Publications Inc.; 2003.

Indzhykulian AA, Stepanyan R, Nelina A, Spinelli KJ, Ahmed ZM, Belyantseva IA, Friedman TB, Barr-Gillespie PG, Frolenkov GI. Molecular remodeling of tip links underlies mechanosensory regeneration in auditory hair cells. PLoS biology. 2013 Jun; 11(6):e1001583. doi: 10.1371/journal.pbio.1001583.

Katoh K, Hammar K, Smith PJ, Oldenbourg R. Birefringence imaging directly reveals architectural dynamics of filamentous actin in living growth cones. Molecular Biology of the Cell. 1999 Jan; 10(1):197–210. doi: 10.1091/mbc.10.1.197.

Kohls O, Holst G, Kühl M. Micro-optodes: The role of fibre tip geometry for sensor performance. SPIE Proc. 1998 Jun; 3483. doi: 10.1117/12.309651.

Kou Jl, Feng J, Ye L, Xu F, Lu Yq. Miniaturized fiber taper reflective interferometer for high temperature measurement. Optics Express. 2010 Jun; 18(13):14245–14250. https://www.osapublishing.org/oe/abstract.cfm?uri=oe-18-13-14245, doi: 10.1364/OE.18.014245.

Liao CR, Hu TY, Wang DN. Optical fiber Fabry-Perot interferometer cavity fabricated by femtosecond laser micromachining and fusion splicing for refractive index sensing. Optics Express. 2012 Sep; 20(20):22813–22818. https://www.osapublishing.org/oe/abstract.cfm?uri=oe-20-20-22813, doi: 10.1364/OE.20.022813.

Liberale C, Cojoc G, Candeloro P, Das G, Gentile F, Angelis FD, Fabrizio ED. Micro-Optics Fabrication on Top of Optical Fibers Using Two-Photon Lithography. IEEE Photonics Technology Letters. 2010 Apr; 22(7):474–476. doi: 10.1109/LPT.2010.2040986.

Martin P, Bozovic D, Choe Y, Hudspeth AJ. Spontaneous oscillation by hair bundles of the bullfrog’s sacculus. The Journal of neuroscience: the official journal of the Society for Neuroscience. 2003 Jun; 23(11):4533–4548.

Nam JH, Peng AW, Ricci AJ. Underestimated Sensitivity of Mammalian Cochlear Hair Cells Due to Splay between Stereociliary Columns. Biophysical Journal. 2015 Jun; 108(11):2633–2647. doi: 10.1016/j.bpj.2015.04.028.

Ozcan Le, Treanton V, Guay F, Kashyap R. Highly Symmetric Optical Fiber Tapers Fabricated With a CO_2_$ Laser. IEEE Photonics Technology Letters. 2007 May; 19(9):656–658. doi: 10.1109/LPT.2007.894963.

Paschotta R. Encyclopedia of Laser Physics and Technology, 2 Volume. Wiley; 2010. https://www.wiley.com/en-gb/Encyclopedia+of+Laser+Physics+and+Technology%2C+2+Volume+Set-p-9783527408283.

Ricci AJ, Crawford AC, Fettiplace R. Active hair bundle motion linked to fast transducer adaptation in auditory hair cells. The Journal of neuroscience: the official journal of the Society for Neuroscience. 2000 Oct; 20(19):7131–7142.

Tobin M, Chaiyasitdhi A, Michel V, Michalski N, Martin P. Stiffness and tension gradients of the hair cell’s tip-link complex in the mammalian cochlea. eLife. 2019; 8. doi: 10.7554/eLife.43473.

Wei T, Han Y, Li Y, Tsai HL, Xiao H. Temperature-insensitive miniaturized fiber inline Fabry-Perot interferometer for highly sensitive refractive index measurement. Optics Express. 2008 Apr; 16(8):5764–5769. https://www.osapublishing.org/oe/abstract.cfm?uri=oe-16-8-5764, doi: 10.1364/OE.16.005764.

Wei T, Han Y, Tsai HL, Xiao H. Miniaturized fiber inline Fabry-Perot interferometer fabricated with a femtosecond laser. Optics Letters. 2008 Mar; 33(6):536–538. https://www.osapublishing.org/ol/abstract.cfm?uri=ol-33-6-536, doi: 10.1364/OL.33.000536.

Yuan L, Wei T, Han Q, Wang H, Huang J, Jiang L, Xiao H. Fiber inline Michelson interferometer fabricated by a femtosecond laser. Optics Letters. 2012 Nov; 37(21):4489–4491. https://www.osapublishing.org/ol/abstract.cfm?uri=ol-37-21-4489, doi: 10.1364/OL.37.004489.

Yuan W, Wang F, Savenko A, Petersen DH, Bang O. Note: Optical fiber milled by focused ion beam and its application for Fabry-Pérot refractive index sensor. The Review of Scientific Instruments. 2011 Jul; 82(7):076103. doi: 10.1063/1.3608111.

